# MitoEdit: a pipeline for optimizing mtDNA base editing and predicting bystander effects

**DOI:** 10.1101/2025.01.22.634390

**Authors:** Devansh Shah, Kelly McCastlain, Ti-Cheng Chang, Gang Wu, Mondira Kundu

**Affiliations:** Department of Cell & Molecular Biology, St. Jude Children’s Research Hospital, Memphis, TN 38105, USA; Center for Applied Bioinformatics, St. Jude Children’s Research Hospital, Memphis, TN 38105, USA; Department of Bioscience and Bioengineering, Indian Institute of Technology Jodhpur, Jodhpur 342030, India

## Abstract

**Motivation:** Human mitochondrial DNA (mtDNA) mutations are causally implicated in maternally inherited mitochondrial respiratory disorders; however, the role of somatic mtDNA mutations in both late-onset chronic diseases and cancer remains less clear. Although recent advances in mtDNA base editing technology have the potential to model and characterize many of these mutations, current editing approaches are complicated by the potential for multiple unintentional edits (bystanders) that are only identifiable through empirical ‘trial and error’, thereby sacrificing valuable time and effort towards suboptimal construct development.

**Results:** We developed MitoEdit, a novel tool that incorporates empirical base editor patterns to facilitate identification of optimal target windows and potential bystander edits. MitoEdit allows users to input DNA sequences in a text-based format, specifying the target base position and its desired modification. The program generates a list of candidate target windows with a predicted number of bystander edits and their functional impact, along with flanking nucleotide sequences designed to bind TALE (transcription activator-like effectors) array proteins. *In silico* evaluations indicate that MitoEdit can predict the majority of bystander edits, thereby reducing the number of constructs that need to be tested empirically. To the best of our knowledge, MitoEdit is the first tool to automate prediction of base edits.

**Availability and implementation:** MitoEdit is freely available at Kundu Lab GitHub (https://github.com/Kundu-Lab/mitoedit).

**Contact:** Corresponding

**Supplementary information:** Supplementary data are available at *Bioinformatics* online.

## Introduction

Maternally inherited primary mitochondrial disorders are caused by pathogenic mutations in the mitochondrial genome (mtDNA) (Ryzhkova, et al., 2018). While somatic mtDNA variants are associated with certain late-onset chronic diseases (e.g., affecting neuromuscular, cardiovascular, and endocrine systems) and cancer (Gorelick, et al., 2021; Marra, et al., 2021; Taylor and Turnbull, 2005), a causal role for most of these variants has yet to be established due to difficulties in engineering relevant models to assess the impact on mitochondrial respiration and other key functions (Gammage, et al., 2018; Lim, 2024; McCormick, et al., 2020; Slone and Huang, 2020). Moreover, the functional impact of most somatic mtDNA variants is inferred by algorithms that consider factors such as population frequency, gene location, sequence conservation, and predicted effect on protein structure (Lott, et al., 2013; Sonney, et al., 2017).

Recent advances in targeted mtDNA base editing have the potential to accelerate the modeling of diseases associated with mtDNA mutations, facilitate preclinical drug testing, and enable therapeutic approaches to correct pathogenic mtDNA mutations (Kar, et al., 2023; Silva-Pinheiro and Minczuk, 2022). MtDNA base editors such as DdCBEs (DddA-derived cytosine base editors) (Mok, et al., 2020; Mok, et al., 2022) and TALEDs (Transcription activator-like effector (TALE)-linked deaminases) (Cho, et al., 2022; Cho, et al., 2024), have demonstrated utility in generating mtDNA mutations across multiple animal and plant models (Chen, et al., 2022; Cho, et al., 2022; Guo, et al., 2022; Guo, et al., 2021; Kang, et al., 2021; Kotrys, et al., 2024; Mok, et al., 2020; Qi, et al., 2021; Sabharwal, et al., 2021). DdCBEs enable C•G-to-T•A conversions through a split-DddA protein that is guided to either side of target DNA by flanking, programmable TALE proteins, each fused to uracil glycosylase inhibitor (UGI) domains and a mitochondrial target sequence (MTS) (Mok, et al., 2020). TALEDs have a similar architecture but enable A•T-to-G•C conversions by replacing UGI domains with a TadA8e domain that is present on either side of the DddA split (Cho, et al., 2022).

Despite potential for disease modeling and therapeutic application, base editors may also introduce additional unintended edits to DNA sequences (Kar, et al., 2023; Wei, et al., 2022). Off-target edits (*i.e*., those that occur outside the target window) may result from either non-specific TALE-DNA interactions or spontaneous reassembly of the split DddA domains (Lee, et al., 2023; Mok, et al., 2020). By contrast, bystander edits may result from the deamination of bases within the target window itself, often as a result of imprecise TALE binding and/or poorly characterized base editing behavior. Current approaches to mitigate unintended edits include designing base editor variants with enhanced DNA-binding domain specificity (Castillo, et al., 2024; Xie, et al., 2024), optimized deaminase activity (Cho, et al., 2024; Huang, et al., 2023), or *in vitro* transcript delivery (Cho, et al., 2024; Guo, et al., 2022). To complement these efforts, we present MitoEdit, an innovative computational tool that identifies optimal windows to facilitate editing of intended target bases while minimizing bystander edits. The tool also predicts the potential impact of unintended bystander edits.

## Tool Description and Software Implementation

MitoEdit is a command-line workflow designed to identify candidate target windows and predict the number and positions of potential bystander edits for each window, based on editing patterns observed in seminal publications (Cho, et al., 2022; Mok, et al., 2020; Mok, et al., 2022). The output includes optimal TALE binding sequences for each window, when such sequences are available. Users input the target base position and desired modification; if the base is targetable, the output is generated as an Excel file with two spreadsheets: the first, detailing the target windows and the second, summarizing the inferred functional consequences of predicted bystander edits (Supplementary Fig. 1A).

This position-based input workflow, along with the output that details the predicted impact of bystander edits, utilizes the human mitochondrial sequence (NC_012920.1) as a reference (Andrews, et al., 1999). Alternatively, users can upload their own DNA sequences as the input file prior to specifying the target base position and the desired modification to facilitate mtDNA targeting in other species or for targeting nuclear DNA. In these contexts, however, the inferred functional impact of bystander edits is not available.

For desired C•G-to-T•A conversions, the canonical cytidine deaminase DddA in DdCBEs has been observed to target cytosines in a *5’-TC* context on either strand (Mok, et al., 2020). The length of the target window ranges from 14 to 18 base-pairs (bp) long, with the position of the target base in the window varying depending on the split site (positions G1397 or G1333) of the enzymatic DddA protein (Fig. 1A-B and Supplementary Table 1A). The workflow generates candidate windows for each split, with the name of each split delineated in the ‘Pipeline’ column of the output file. Additionally, the workflow accommodates pipelines for the improved *5’-HC* context (H=A, C, or T) based on the evolved DddA variants (Mok, et al., 2022) (Supplementary Fig. 1B and Supplementary Table 1A). Since TALEDs also utilize the DddA domain, MitoEdit assumes that the target base should occur adjacent to a cytosine on either strand (Yin, et al., 2023). Thus, for desired A•T-to-G•C conversions, editing parameters similar to DdCBEs are observed with the exception that the target base must be an adenine in a *5’-SA* or *5’-AS* context (S = G, C) to accommodate the TadA-derived deoxyadenosine deaminase domain (AD) (Supplementary Fig. 1C and Supplementary Table 1A). Notably, while multiple TALED designs have been described (Cho, et al., 2022), MitoEdit only models the split TALED (sTALED) configuration with the AD fused to either TALE. This is because the sTALEDs share a similar architecture with DdCBEs and exhibit higher average editing efficiency compared to other TALEDs.

**Figure 1.**
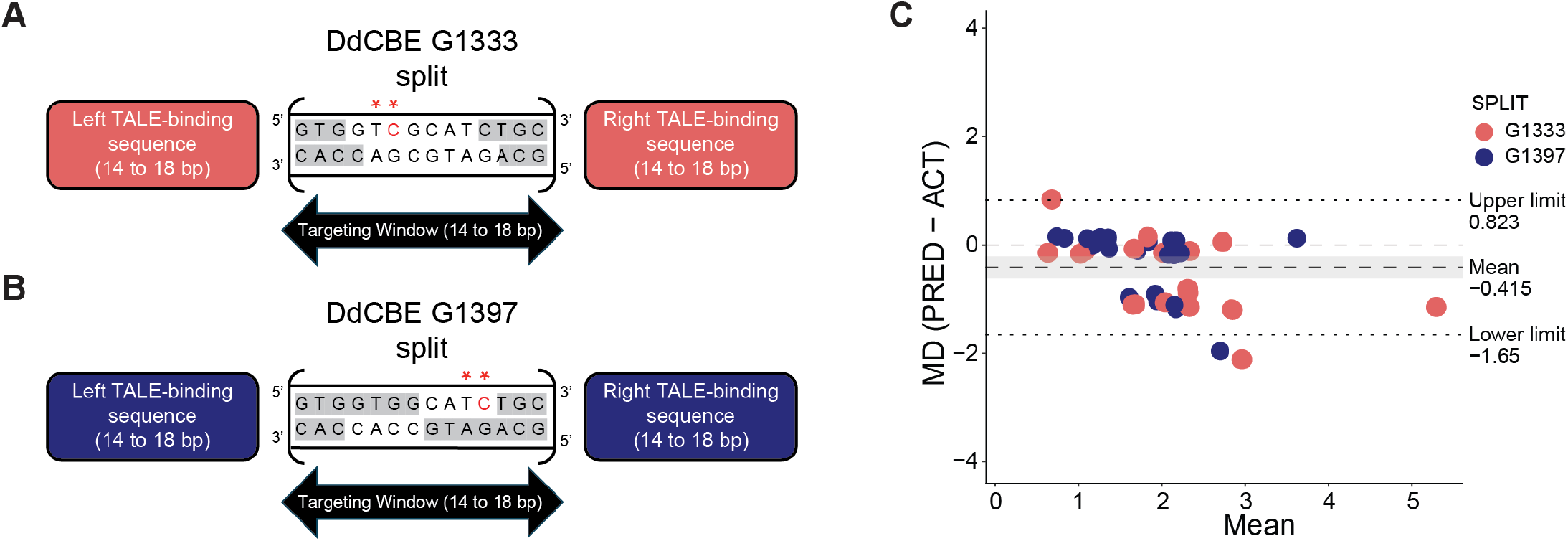
MitoEdit pipelines and validation. (A, B) Diagrams depicting the editing patterns observed by DdCBE base editors among select publications used for validation. Cytosines within a *5’-TC* context (double red asterisks) are edited by the G1333 DddA split (A) when located 4-10 bp from the 5’ end (A) and are edited by the G1397 DddA split (B) when located 4-7bp from the 3’ end (Mok, et al., 2020). Target cytosines (red font) within the regions highlighted in white are accessible to the base editor for C>T editing. (C) Bland-Altman plot illustrates concordance between edits predicted by MitoEdit and those observed (n = 41) across the publications used for validation. The grey ribbon highlights the 95% confidence interval for the mean difference (MD) between measures versus no difference (MD, -0.4151; P < 0.001, 1-sample t-test). Dashed lines indicate the upper and lower limits for the mean difference. Abbreviations: PRED, predicted; ACT, actual.

The MitoEdit workflow utilizes the TALE-NT 2.0 tool to select optimal TALE binding sequences for each candidate target window (Doyle, et al., 2012). Lengths for TALE array repeat variable diresidues (RVDs) range from 14 to 18 bp. If matching flanking TALE arrays are not found, users can easily adjust the TALE-NT input to explore additional options. For bystander edits that occur within human mtDNA (NC_012920.1), the predicted functional impact is called using a previously described pipeline (McCastlain, et al., 2024); however, only the impact of single-base edits is inferred. When target and bystander edits impact a single codon, MitoEdit flags (but does not interpret) the result.

The potential utility of MitoEdit in streamlining construct design is highlighted using the m.11696G>A (*MT-ND4)* mutation, which is associated with Leber’s hereditary optic neuropathy (LHON). For this example, the command *python mitocraft.py 11696 G A* is executed which instructs MitoEdit to read the default human mtDNA file, to target bp position 11,696 with a G>A conversion, and to output the relevant information (Supplementary Table 1B,1C). Here, MitoEdit predicts that the m.11696G>A mutation can be introduced with windows containing a single bystander edit at positions 11698 or 11704, both of which result in synonymous mutations. Additional detailed instructions and tutorials pertaining to MitoEdit are freely available under an MIT License through GitHub (https://github.com/Kundu-Lab/mitoedit).

## Validation and Application

The MitoEdit workflow was validated by comparing the number of edits predicted by the pipeline to the actual number of edits reported by multiple studies (Chen, et al., 2022; Guo, et al., 2022; Guo, et al., 2021; Kang, et al., 2021; Lee, et al., 2021; Mok, et al., 2020; Mok, et al., 2022; Qi, et al., 2021; Sabharwal, et al., 2021). Using a Bland-Altman plot to assess agreement between measures (Fig. 1C), we found that the mean difference between predicted and actual edits was close to zero for 59% (24/41) of the data points, resulting in a Lin’s concordance correlation coefficient of 0.651 (95% CI: 0.306,0.807). A modest departure between measures was noted when three or more edits were observed, resulting from underestimation of actual edits (mean difference: -0.415; 95% CI: -0.614, -0.215). This discrepancy is most likely attributable to sequencing errors, misreporting background heteroplasmies as low-level base edit conversions, or previously uncharacterized rules governing base editing behavior. Towards the latter, we noted multiple cases where C>T editing occurred in *5’-AC* contexts within the 4-10 bp window (G1333 split only). For both splits, we observed C>T editing in *5’-TC* contexts when the thymine (G1333) or cytosine (G1397) was just outside of the editing window.

MitoEdit’s strength resides in its ability to quickly assess the feasibility of engineering mutations from extensive lists of targets. To demonstrate this point, we queried the MITOMAP database (Lott, et al., 2013) to identify targetable mutations and editing windows that yielded the least number of bystander edits (preferably benign) (Supplementary Table 1D). Of the 91 confirmed pathogenic or likely pathogenic mutations catalogued, the pipeline identified 54 (59.3%) point mutations as viable targets (Supplementary Table 1E-). MitoEdit predicted that one-third (n=18) of these targets could be edited without any bystander edits. An additional 18 could be targeted with one bystander edit; however, of these, only three were predicted to be benign (Supplementary Table 1E).

## Conclusion

MitoEdit is a novel pipeline designed to identify optimal target windows for engineering mutations into the mitochondrial genome with DddA-derived base editors. Optimization includes the ability to identify candidate windows with the least number of bystander edits and to pair these windows with flanking TALE-binding sequences. Utility is also demonstrated through our candidate window analysis of all confirmed point mutations currently listed in MITOMAP. Additionally, as deep-sequencing data from future studies becomes publicly available, and as base editors continue to evolve, MitoEdit can be adapted to integrate machine learning models to identify more complex editing patterns and to predict editing efficiencies for target and bystander edits.

---

**Supplementary Figure 1.**
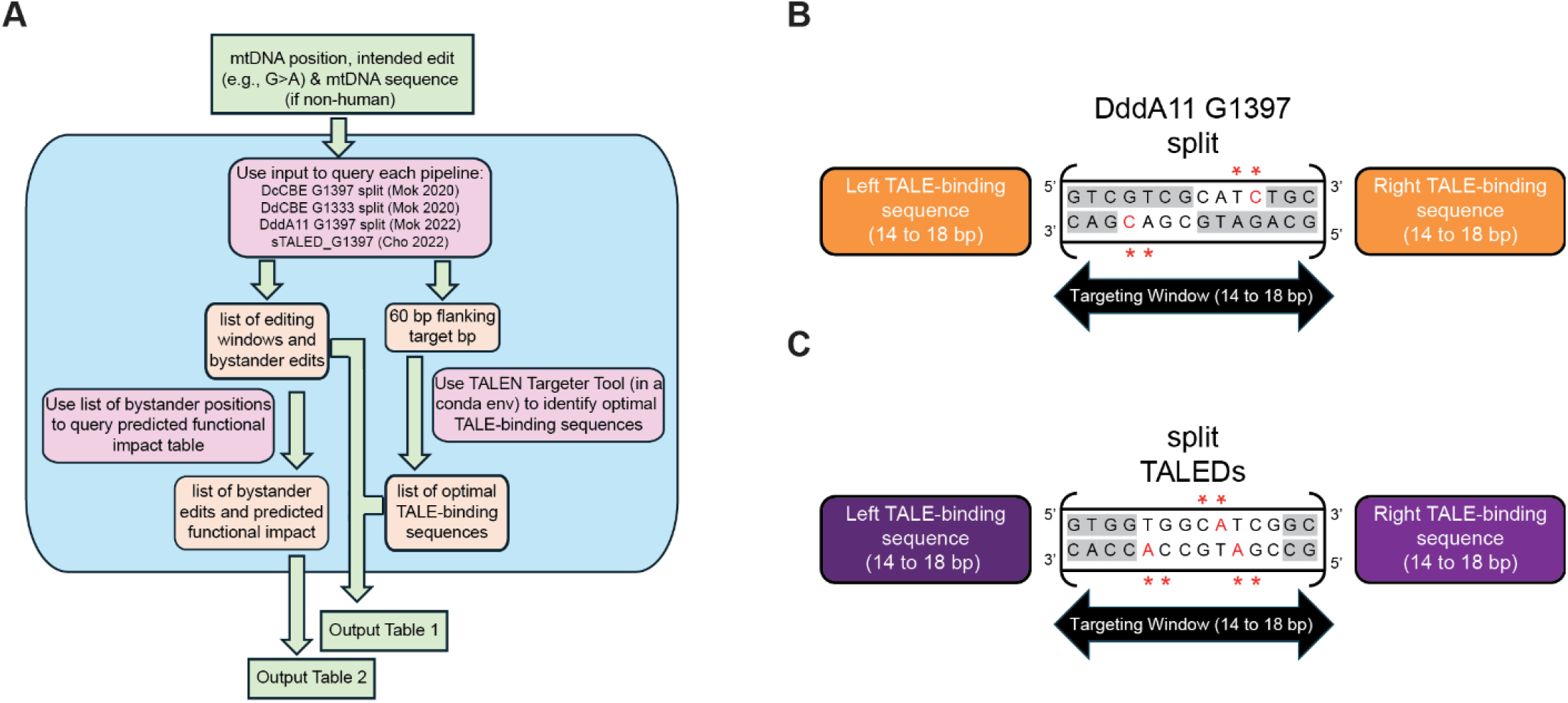
MitoEdit pipelines for evolved DddA variant and split TALEDs. (A) MitoEdit pipeline workflow. (B) The evolved DddA11 variant exhibits a similar pattern to the DdCBE G1397 split, but with a modified editing context (*5’-HC*, where H = A, C or T) to target cytosines. (C) The split TALED (sTALED) base editor targets adenines 5-12 base pairs (bp) from the end closest to the TALE (right or left) fused to the deoxyadenine deaminase (AD) domain. Here, the left-binding TALE sequence is depicted in dark purple, to indicate where the TALE protein that is fused to the AD will bind. sTALEDs follow an editing pattern similar to the DdCBE G1397 split, but with a modified context (*5’-SA* or *5’-AS*, where S = G or C) to target adenines. In both (B) and (C), regions highlighted in white are accessible to the base editors. Double red asterisks indicate the editing context and red font indicates the targeted base(s).

